# Mammalian heat shock protein A4 family ortholog *Hsc70Cb* is required for two phases of spermatogenesis in *D. melanogaster*

**DOI:** 10.1101/2024.12.09.627449

**Authors:** Brendan J. Houston, Joseph Nguyen, Richard Burke, Andre N. Alves, Gary Hime, Moira K. O’Bryan

## Abstract

Heat shock proteins play essential roles as molecular chaperones, enacting protein folding and preventing of protein aggregation. In a previous study, a predicted damaging homozygous non-synonymous genetic variant was detected in the heat shock protein gene *HSPA4L*. Here, we used RNA interference in *Drosophila melanogaster* to explore the role of the heat shock protein A member 4 family (HSPA4) family in male fertility. Expression of the fly orthologue of the mammalian *HSPA4* and *HSPA4L* genes, *Hsc70Cb*, was ablated in the male germline using two RNAi lines and the Nanos-Gal4 driver. Strong knockdown of *Hsc70Cb* in male germ cells resulted in male sterility, characterised by the absence of germ cells in testes and the over-proliferation of the testis soma. A less robust knockdown of *Hsc70Cb* in the male germline resulted in a sperm individualisation defect and a failure of sperm release into the seminal vesicle (analogous to the epididymis). When human *HSPA4* or *HSPA4L* cDNA was introduced into infertile *Hsc70Cb* mutant flies, a partial rescue was observed, whereby in the robust *Hsc70Cb* knockdown setting germ cells progressed to the spermatocyte stage before undergoing cell death. Collectively, the absence of sperm in the *Hsc70Cb* (line 1) is consistent with the infertile man harbouring a homozygous *HSPA4L* genetic variant, supporting the hypothesis that *HSPA4L* is required for male fertility in humans and flies and highlighting the utility of the fly as a model of human spermatogenesis.

**In brief:** *Hsc70Cb* is essential for spermatogonia survival and sperm individualisation in *Drosophila*. This study highlights the conserved roles of the HSPA4 family across animals and the utility of flies as a model organism for male fertility research.

## Introduction

Male infertility is a prevalent condition affecting more than 7% of men worldwide and is caused by a myriad of genetic, lifestyle and other factors (Houston et al., 2021c, Kimmins et al., 2024). Owing to the sheer number of genes expressed in the testis in mammals (>15,000 in men), there are numerous opportunities for genetic variants to impact male fertility (Djureinovic et al., 2014, Soumillon et al., 2013, Houston et al., 2021a). In support of a strong genetic component, we and others have used next generation sequencing to identify a number of likely genetic causes of male infertility (Oud et al., 2022, Nagirnaja et al., 2022, Oud et al., 2021, Oud et al., 2020), many of which require further validation. Animal models are vital for such validation, notably when replication studies in humans are yet to identify recurrent variants in the affected genes. While mouse models are the gold standard for validation studies, fly (*Drosophila*) models have also shown considerable utility for studying human male fertility defects (Netherton et al., 2024, Ayers et al., 2023, Yu et al., 2015, Sieper et al., 2024) and notably 75% of disease-causing genes are conserved in flies (Ugur et al., 2016).

In a previous study, a rare and predicted damaging homozygous non-synonymous genetic variant was detected in the heat shock protein gene *HSPA4L* (Chr4:127827312 G>A (hg38)) in a man with non-obstructive azoospermia (Nagirnaja et al., 2022). The variant has a high pathogenicity score (CADD = 28.2, *P* = 7.178562 x 10^-6^) and was predicted to be disease causing by MutationTaster. Heat shock proteins play essential roles in governing protein folding and the prevention of protein aggregation to balance protein homeostasis, ‘proteostasis’ (Saibil, 2013). In performing this role, heat shock proteins play critical roles in the initial folding of nascent polypeptides and the re-folding of proteins that have become denatured, as well as in protein trafficking and in the coordination degradation of proteins tagged for destruction (Hartl et al., 2011). Consistent with the thousands of proteins expressed during spermatogenesis (Djureinovic et al., 2014, Uhlen et al., 2015), heat shock proteins are vital to ensure proteostasis in germ cells for successful sperm production (Nixon et al., 2017).

Knockout mouse studies identified that the ablation of *Hspa4l* rendered 42% of males infertile (Held et al., 2006) and similarly the deletion of gene family member *Hspa4* caused infertility in 61% of males (Held et al., 2011). The loss of *Hspa4* caused significant germ cell apoptosis during meiosis, a reduction in epididymal sperm count, and resulted in poor sperm motility. Similarly, the absence of *Hspa4l* resulted in very few numbers of sperm in the epididymis and significantly compromised sperm motility (Held et al., 2006). HSPA4L has also been localised to the mammalian sperm midpiece, where it likely contributes to sperm motility, as highlighted by the finding that HSPA4L content is associated with improved sperm motility in men (Liu et al., 2019). Moreover, testis HSPA4L content reduces with age (Liu et al., 2019), potentially contributing to subfertility in older men. While *Hspa4l* is clearly required for optimal male fertility in mice, the phenotype is not a perfect match with the infertile, azoospermic man containing the *HSPA4L* variant. We thus sought to explore *HSPA4L* using flies as a model system.

An analysis of *Drosophila* orthologues for human *HSPA4L* revealed a single gene in flies, *Hsc70Cb*, which was predicted to be a conserved ortholog of both gene family members *HSPA4* and *HSPA4L*. We therefore aimed to further investigate the role of *HSPA4* gene family function in male fertility using testis-specific knockdown fly models, targeting the fly ortholog *Hsc70Cb* with commercially available RNAi lines.

## Materials and methods

### Analysis of Hsc70Cb expression

Data on *Hsc70Cb* expression were extracted from an existing fly testis single RNA sequencing library (Witt et al., 2019), containing clusters for somatic hub and cyst cells, and germ cells including early and late spermatogonia, spermatocyte and spermatids.

### Orthologue comparison

The Drosophila RNAi Screening Center Integrative Ortholog Prediction Tool was used to identify *Drosophila* orthologues for the human *HSPA4L* gene. The highest ranked ortholog for *HSPA4L* was *Hsc70Cb*, which is also the highest ranked hit for mammalian gene family member *HSPA4*, suggesting gene duplication events may have arisen in mammals to produce two closely related genes. This study therefore investigates the function of the HSPA4 family in male fertility in flies.

### Drosophila models

Commercially available fly stocks encoding for *Hsc70Cb* RNA interference (RNAi) were purchased from the Vienna *Drosophila* Resource Center, Austria (VDRC) and the Bloomington *Drosophila* Stock Center, USA (BDSC), as shown in Table 1. The *w^1118^* line was used as a wild type control, which was crossed to the Gal4 driver line used for each experiment (Table 1) to generate a background matched control. In order to induce testis specific RNAi, we utilised the germ-cell specific *Nanos*-Gal4 (Van Doren et al., 1998). Specifically, a Nanos-Gal4 strain containing a UAS-Gal4 element was used to improve knockdown efficiency (Netherton et al., 2024). For testis somatic cell knockdown, we used the Traffic jam-Gal4 line (Li et al., 2003). Flies were maintained in plastic vials on standard medium (Richards and Burke, 2015) at 22-25 °C until used for crossing. Crossing was undertaken at 29 °C to maximise Gal4 activity unless otherwise stated.

**Table 1.** Fly lines used in this study, sourced from Bloomington (BDSC), Vienna (VDRC) or Kyoto (DRRG) *Drosophila* centres.

| Line | Source | Function |
| --- | --- | --- |
| $w^{1118}$ (B3605) | BDSC, 3605 | Wild type control |
| *Nanos*-Gal4; UAS-Gal4 | Generated by crossing BDSC 4937 and DRRG 108492 | Spermatogonia and early spermatocyte Gal4 driver |
| *Nanos*-Gal4; don juan-GFP | Generated by crossing BDSC 4937 and BDSC 58406 | Spermatogonia and early spermatocyte Gal4 driver; sperm tail-GFP |
| *Traffic jam*-Gal4 | DRRG 104055 | Cyst stem cell, early cyst cell Gal4 driver |
| *Act5c*-Gal4 (Actin) | BDSC 3953 | Whole body Gal4 driver |
| *Hsc70Cb* knockdown 1 | VDRG 330137 | *Hsc70Cb* RNAi line 1 |
| *Hsc70Cb* knockdown 2 | BDSC 33742 | *Hsc70Cb* RNAi line 2 |
| *HSPA4* human cDNA | DRRG 306061 | Human *HSPA4* for rescue cross |
| *HSPA4L* human cDNA | In house | Human *HSPA4L* for rescue cross |

### Fertility assays

Males from *Hsc70Cb* RNAi stocks and the control *w^1118^* line were crossed to virgin females from *Nanos*-Gal4; UAS-Gal4 or *Traffic jam*-Gal4 drivers at 29 °C. Females were allowed to lay progeny for one week, before all parent flies were removed from the vials. The resulting F_1_ males were collected and aged to 2-5 days post-eclosion from pupal cases. Each male was then individually housed with two *w^1118^* virgin females for 48 h at 29 °C to assess fertility. Adult flies were then removed, and the total number of pupae generated from each breeding test counted per vial 5 days later. This experiment was then repeated, using males raised at 22 and 25 °C to dissect the impact of varying levels of RNAi knockdown wherein high ambient temperatures lead to improved knockdown efficiency. For the second experiment, males were fertility tested as described above. Pupae numbers were counted after 8 and 10 days for the 25 and 22 °C conditions, respectively, noting that speed of developmental timing is determined by environmental temperature.

### Assessment of the male reproductive tract

Testes, seminal vesicles, and accessory reproductive glands were dissected from 2-4-day old adult males generated from the *Hsc70Cb* RNAi x *Nanos*-Gal4; UAS-Gal4 crosses in a drop of PBS on a glass slide. Testes and seminal vesicles were separated from the ejaculatory duct and accessory glands, except for *Hsc70cb* RNAi 1 line, where ejaculatory ducts were left joined to the seminal vesicles due to the tiny testis size. In other experiments, larval male gonads were isolated from third instar larvae to inspect juvenile testis histology. Histology was observed using both brightfield and phase contrast microscopy, as described previously (Netherton et al., 2024).

To perform immunostaining, whole testes were fixed in 4% paraformaldehyde for 30 min at room temperature, then washed in 1 × PBS and permeabilised in 0.2% Triton-X-100 (Sigma Aldrich) in PBS for 30 min. Testes were then washed in 1 × PBS and stained with markers to investigate spermatogenesis (Table 2). To do so, testes were blocked in CAS-block (Agilent, Santa Clara, CA, USA) for 30 min then incubated overnight at 4 °C in primary antibody diluted in Dako antibody diluent (Agilent) at concentrations shown in Table 2. After washing in 1 × PBS, testes were incubated in secondary antibody (Table 2) for 1 h at room temperature and then washed in 1 × PBS. Testes were counterstained with 10 μg/ml DAPI nuclear stain (ThermoFisher Scientific, Waltham, MA, USA) for 30 min, washed in 1 × PBS and whole mounted on slides with Dako Fluorescence Mounting Media (Agilent). Similarly, both RNAi lines were also crossed to a *Nanos*-Gal4; don juan-GFP line to define if spermatids were present in mutant testes. Testes and seminal vesicles were processed as above. Males generated from the *Nanos*-Gal4; UAS-Gal4; don juan-GFP cross were also allowed to mate with *w^1118^* females for 3 days. Afterwards, the spermathecae and seminal receptacles were dissected to test whether sperm from mutant males could be found within the female reproductive tract.

**Table 2.**
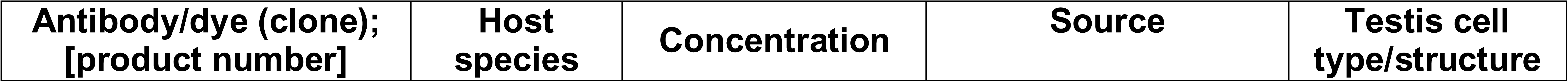
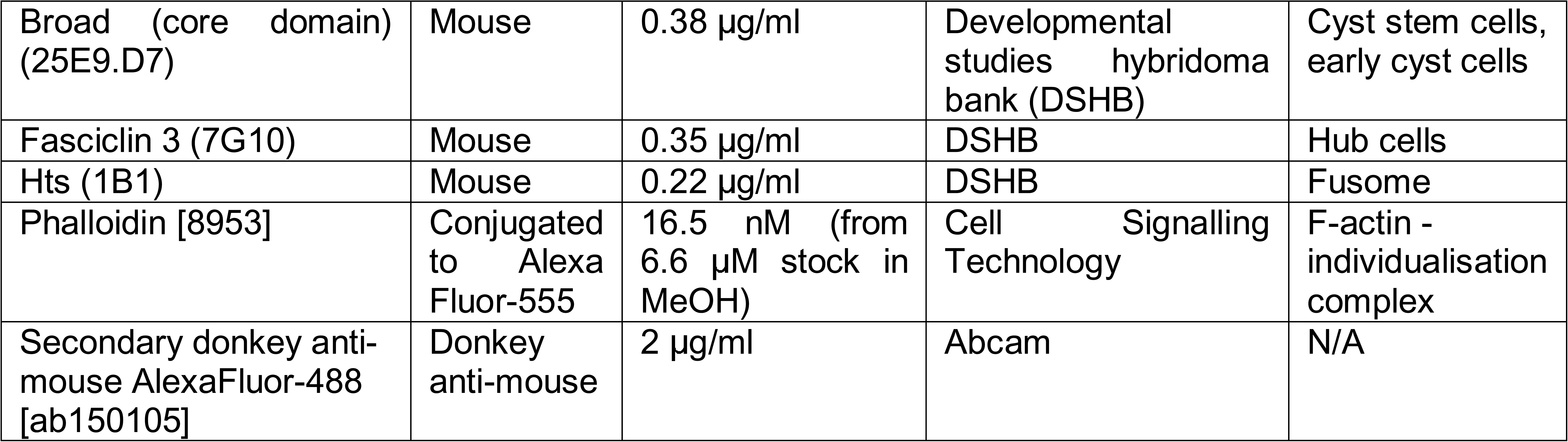
Antibodies and markers of spermatogenesis used.

Ovary histology was similarly investigated using phase contrast and immunofluorescence imaging following staining with DAPI, as described above.

Slides were imaged on an Olympus BX-53 microscope (Olympus America, Center Valley, PA, USA), CV1000 confocal microscope (Olympus America, USA) or Zeiss Axioskop 2 microscope (Zeiss, Germany).

### Human HSPA4 and HSPA4L cDNA rescue

To investigate if human HSPA4 or HSPA4L cDNA could rescue the infertility phenotypes arising from RNAi knockdown in the germline, we generated compound lines containing the fly RNAi, Nanos-Gal4 and human cDNA elements. To do so, we sourced an existing transgenic fly line encoding human UAS-HSPA4 cDNA (Table 1) and generated a line encoding human *HSPA4L* cDNA. For the latter, we cloned *HSPA4L* from human testis cDNA into the pUAST_attB vector using primers F – GCGGTACCATGTCTGTGGTTGGCATTGACCTCGGCTT and R – GCTCTAGATTAGTCCACTTCCATCTCTCCAG. pUAST *HSPA4L* vector was injected it into fertilised *yw, PhiC31; attP86Fb* eggs (i.e., 1 cell stage zygotes). Lines were balanced over the TM6B balancer chromosome. The capacity of human HSPA4 and HSPA4L to individually rescue the infertility phenotype was investigated by fertility testing and histology as detailed above.

### Quantitative PCR

*Hsc70Cb* expression was investigated using quantitative PCR as described previously (Houston et al., 2021b), relative to housekeeper gene *Rpl11*. RNA was extracted from whole male flies generated by crossing each RNAi line to an *Act5c*-Gal4 driver (Table 1), generating a whole-body knockdown. Primer sequences were *Hsc70Cb* F:CGAACTCACGAGCATTCCAG, R:ACGCTCTGCATTGGTAAAGAAC; *Rpl11* F:GGTCCGTTCGTTCGGTATTCGC, R:GGATCGTACTTGATGCCCAGATCG.

### Statistical analysis

GraphPad Prism 10 was used for statistical analysis, wherein a one-way ANOVA with Tukey post-hoc test was conducted to investigate significance of pupae count and mRNA expression data in comparison to the control strain (*w^1118^* on the respective genetic background). Significance was denoted as follows: * *p* <0.05, ** *p* < 0.01, *** *p* < 0.001 and **** *p*< 0.0001.

## Results

### *Hsc70Cb* is enriched in early spermatogonia

Data from a published single cell RNA sequencing library for fly testes (Witt et al., 2019) indicated that *Hsc70Cb* is expressed at low levels in all cell types in the testis but elevated in early spermatogonia (Figure 1).

**Figure 1.**
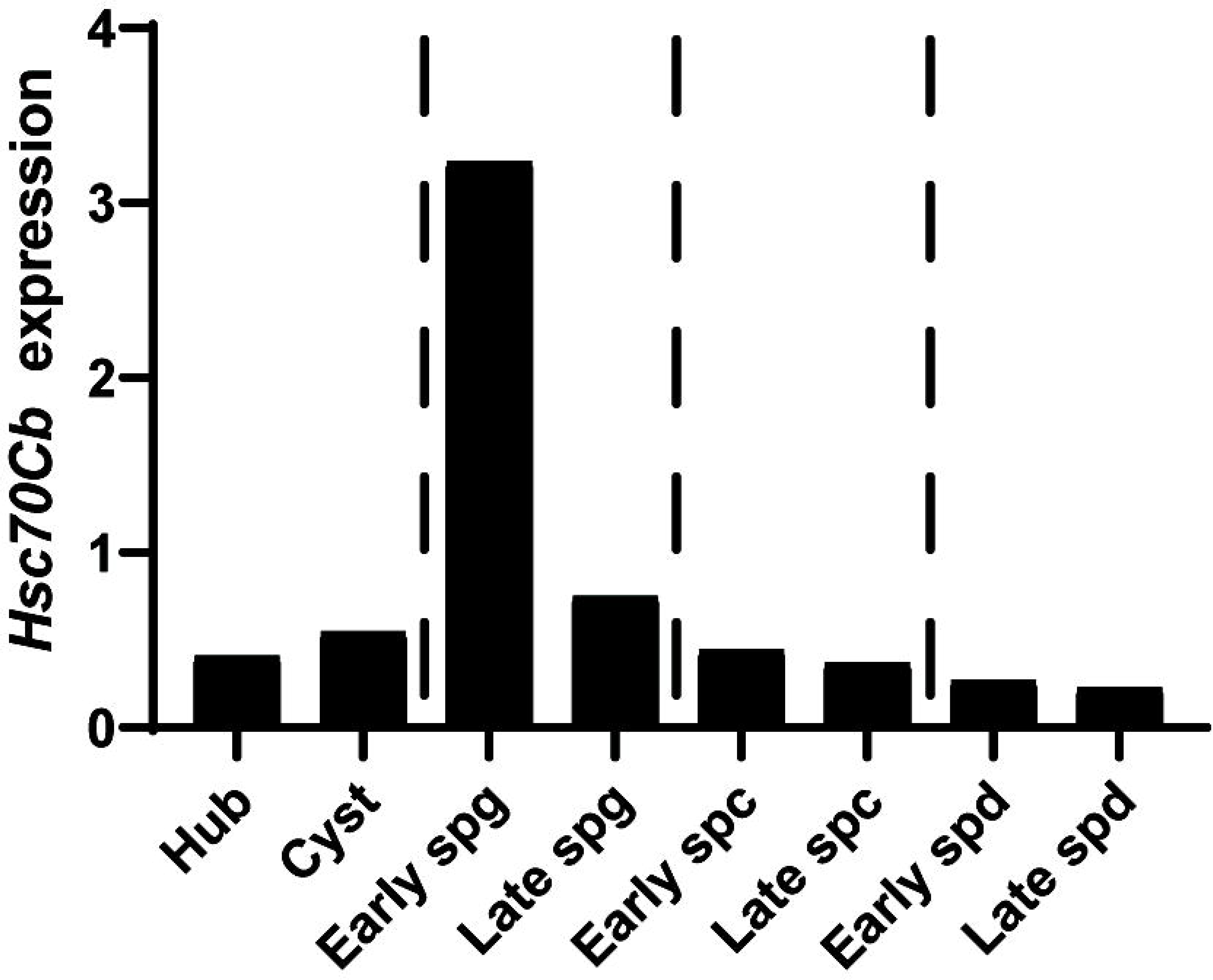
*Hsc70Cb* is expressed primarily in early germ cells in *Drosophila melanogaster* testes. A. *Hsc70Cb* expression in major germ and somatic cell types in fly testes as determined by single cell sequencing data (Witt et al., 2019). Spg = spermatogonia, spc = spermatocyte, sp(t)d = spermatid.

### *Hsc70Cb* is required for male germ cell function but not testis somatic cell function

To investigate the role of *Hsc70Cb* in male fertility, we sourced two RNAi lines targeting *Hsc70Cb* transcripts and crossed these to the Nanos-Gal4 (germ cell-specific) and Traffic-jam Gal4 (testis somatic cell-specific) drivers to induce RNAi knockdown in two major cell lineages of the testis (Figure 2A). Knockdown of *Hsc70Cb* in male germ cells (*Nanos*-Gal4 driver) by both RNAi lines at 29 °C resulted in male sterility (Figure 2A; *p* < 0.0001). When RNAi of *Hsc70Cb* was driven within the somatic cell population of the testis (Traffic jam-Gal4), male fertility was unaffected (Figure 2B; *p* = 0.2). Next, the germ cell specific knockdown of *Hsc70Cb* was repeated over a range of 22-29 °C to determine the effects of temperature on the fertility phenotype (Figure 2C). We observed that the sterility in males generated from the *Hsc70Cb* RNAi 1 line was independent of experiment temperature and was apparent at 22, 25 and 29 °C. *Hsc70Cb* knockdown by RNAi 2 line, however, only induced sterility at 29 °C and males were fertile when reared at 22 and 25 °C, likely attributable to lower Gal4 activity at 22 and 25 °C. Quantification of mRNA levels was next assessed in male flies within which RNAi was driven by *Act5C-Gal4*. This revealed a knockdown efficiency of 84% (*p* < 0.0001) for RNA line 1 and 27% (*p* = 0.0053) for RNA line 2 relative to controls (Figure 2D). A significant difference was also observed between RNA lines 1 and 2 (*p* < 0.0001). Thus, RNAi line 1 represented a strong knockdown and RNA line 2 a weak knockdown for *Hsc70Cb*.

**Figure 2.**
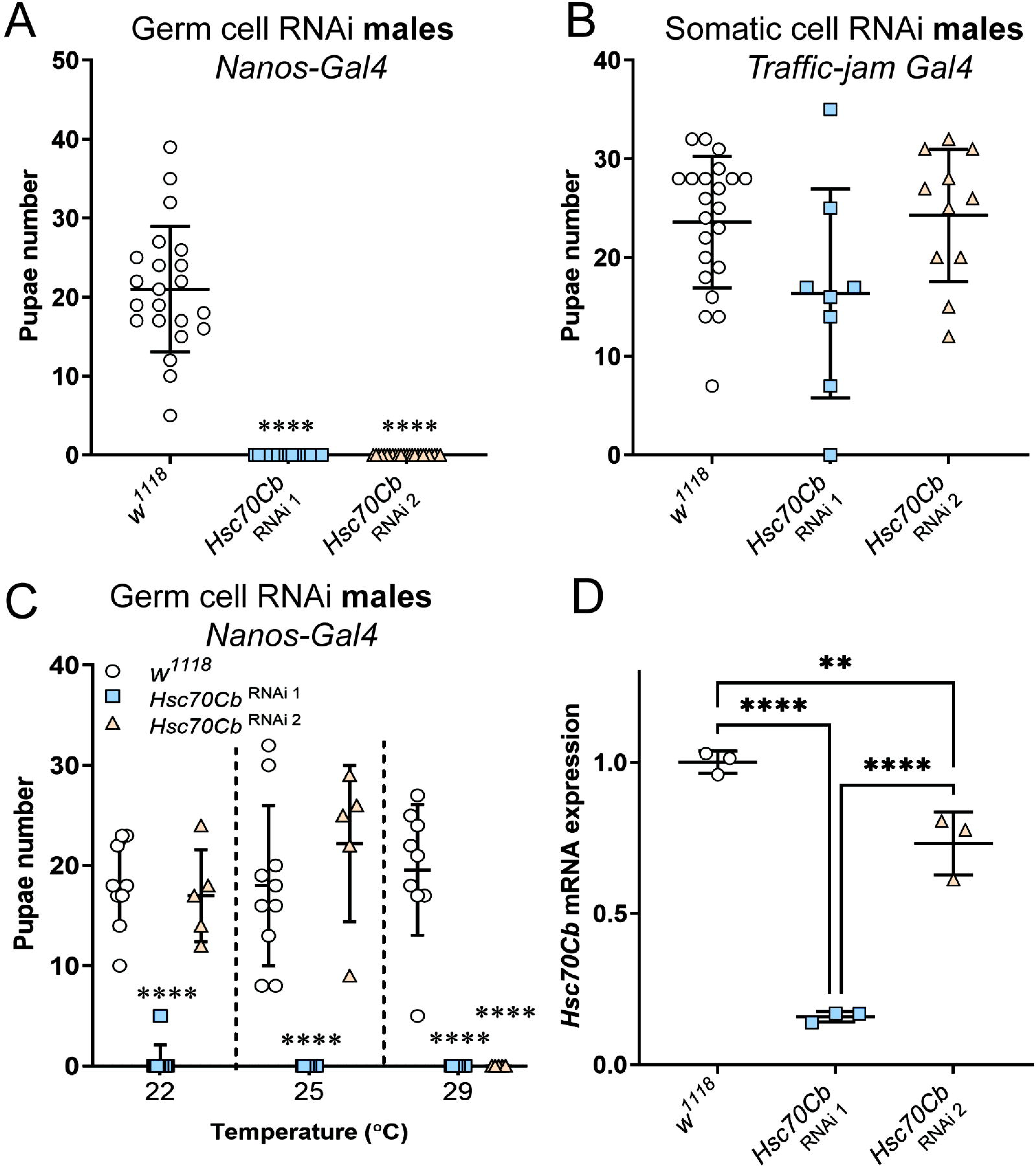
*Hsc70Cb* is required in the germline for male fertility. RNAi knockdown of *Hsc70cb* was performed within the germ cell (A) or somatic cell (B) lineages of the testis, using *Nanos* and *Traffic jam* drivers, respectively. (C) Germ cell RNAi was repeated over a temperature range of 22-29 °C. (D). Quantification of *Hsc70Cb* mRNA levels relative to housekeeper gene *Rpl11*. Whole body knockdown was driven in male flies using the Act5C driver. *w^1118^* (on the same genetic background as RNAi lines) was used a fertile control. **** = *P* < 0.0001 in comparison to *w^1118^*.

Within females (Supplementary Figure 1A), knockdown of *Hsc70Cb* with the RNA 1 line under Nanos-Gal4 control induced sterility (*p* = 0.0004) but females generated by the RNAi line 2 cross were fertile and comparable to *w^1118^* controls (*p* = 0.99).

Collectively, these results suggest the *Hsc70Cb* is required for male and female fertility in flies, with a role in germ cells. This study largely focuses on the requirement of *Hsc70Cb* in male germ cell function.

### *Hsc70Cb* is essential for male germ cell development

To investigate the cause of male infertility in both lines targeting *Hsc70Cb* RNAi, we assessed testis histology in third instar larvae and adults (Figure 3). In *w^1118^* controls, male larval gonadal discs were normal and round (Figure 3A, B). Staining with the fusome marker (Figure 3C), Hts, revealed a normally established fusome interweaved between cysts of spermatogonia and spermatocytes. The fusome is a germline specific organelle that allows communication between sister germ cells and is evidence of intact cysts (Kaufman et al., 2020). Knockdown of *Hsc70Cb* by RNAi 1 line caused a striking reduction in larval testis size (‘tiny testis’; Figure 3D, E) in all males investigated. A large cluster of cells were present at the hub, but fusome staining was not apparent, suggesting early germ cells were absent (Figure 3E, F). The larval testis size in males from the *Hsc70Cb* RNAi 2 line was equivalent to that of *w^1118^* controls (Figure 3G) and observed normal fusome staining in these testes, suggesting normal establishment of spermatogenesis but a difference in the phenotype of the two RNA lines targeting *Hsc70Cb*.

**Figure 3.**
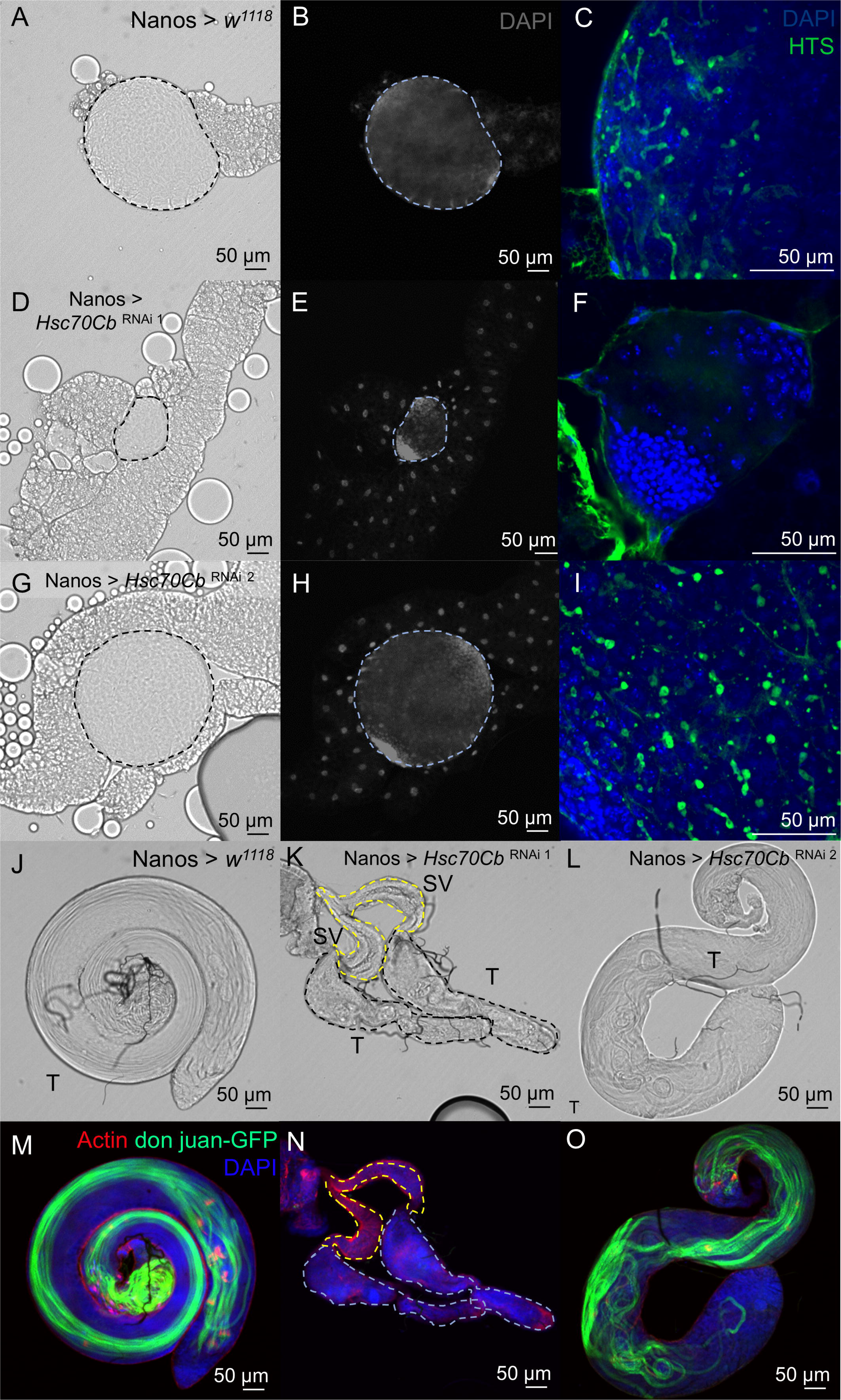
*Hsc70Cb* is essential for normal spermatogenesis and testis morphology. (A-I) Testes were isolated from third instar larvae control (*w^1118^* on a *Nanos*-Gal4; UAS-Gal4 background) and *Hsc70Cb* mutants driven by *Nanos*, stained with DAPI and probed for HTS (fusome marker; green). (A-C) *w^1118^*, (D-F) *Hsc70Cb* RNAi line 1, (G-I) *Hsc70Cb* RNAi line 2. (J-O) Adult testes from a similar cross containing don juan-GFP were stained with DAPI phalloidin (staining F-actin [individualisation complex]; red). (J, M) *w^1118^*, (K, N) *Hsc70Cb* RNAi 1, (L, O) *Hsc70Cb* RNAi 2. Scale bars = 50 µm. T = testis, SV = seminal vesicle

In *w^1118^* adults, testes were of normal size and morphology – coiled tubes (Figure 3J; Supplementary Figure 2A). Spermatids were detected using a *Nanos* driver line containing don juan-GFP protein as a marker of the sperm tail (green; Figure 3M). Co-staining with phalloidin as a marker of actin bundles (red) revealed spermatids were individualising normally. By contrast, testes were abnormally small in males from the *Hsc70Cb* RNAi line 1 (Figure 3K; Supplementary Figure 2B) and lacked don juan-GFP staining (Figure 3N) indicating an absence of elongating spermatids or sperm. While testes from adult males of the *Hsc70Cb* RNAi 2 line were of a normal size, they were histologically abnormal (Figure 3L). Notably, DAPI staining revealed a mass of disorganised spermatids at the distal testis (Supplementary Figure 2C) and spermatid bundle misalignment as marked by don juan-GFP (Figure 3O). The presence of poorly organised spermatids suggested defects in sperm individualisation. By comparison, sperm generated by the *w^1118^* control were neatly organised into bundles spanning the length of the testis (Figure 3M).

We used phase contrast histology to further explore the cellular content of testes from each cross (Figure 4). In *w^1118^* control testes, numerous cysts of developing germ cells and bundles of spermatids were observed (Figure 4A, D). Spermatids were individualised and present within the seminal vesicles (Figure 4A). By contrast, in the *Hsc70Cb* RNAi 1 line there was an almost complete absence of intact cells in the testis (Figure 4B, E). Most cells were amorphous and appeared to be dying. No spermatocytes or spermatids were detected, and the seminal vesicles were devoid of sperm. A similar phenotype was seen in the ovary (Supplementary Figure 1), where ovarioles contained oocytes of various developmental stages in the *w^1118^* control (Supplementary Figure 1B-D, H) but lacked oocytes in RNAi line 1 females (Supplementary Figure 1E-G, I-J).

**Figure 4.**
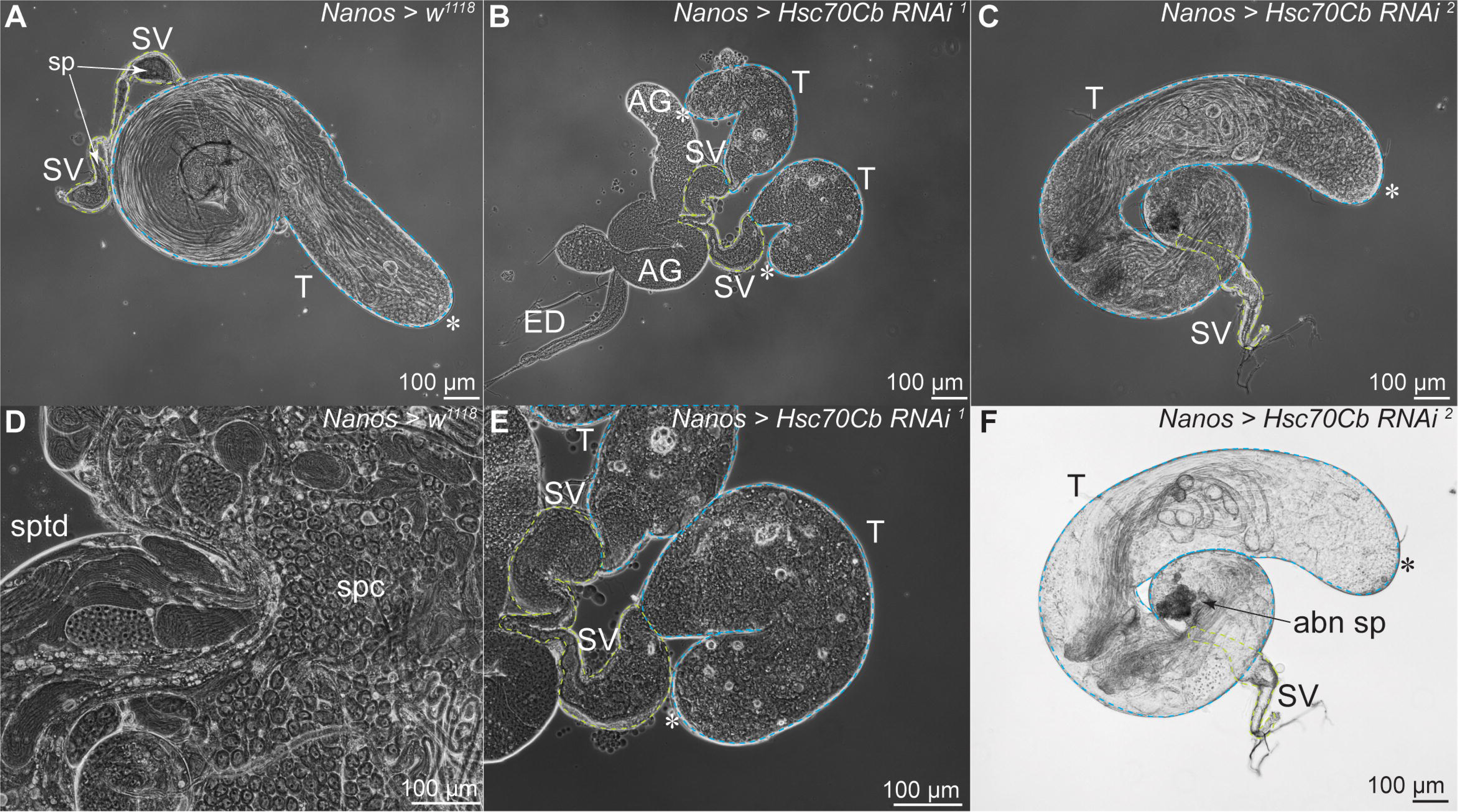
*Hsc70Cb* is required for germ cell survival and sperm individualisation. Phase contrast images of control and *Hsc70Cb* mutant testes. (A, D) *w^1118^*, (B, E) *Hsc70Cb* RNAi 1, (C, F) *Hsc70Cb* RNAi 2. Scale bars = 100 µm. T = testis, SV = seminal vesicle, AG = accessory gland, ED = ejaculatory duct, sp = sperm, spc = spermatocyte, sptd = spermatid, abn = abnormal, * = hub.

As defined above, while spermatids were detected in the testes of *Hsc70Cb* RNAi 2 line males using phase contrast imaging (Figure 4C, F), they were abnormally coiled and undergoing degradation (arrow) due to a predicted failure of sperm individualisation. No sperm were detected in the seminal vesicles (Figure 4C, F). To confirm this finding, testes were stained with DAPI and phalloidin (actin marker) (Figure 5). In *w^1118^* control testes, discrete sperm bundles were observed, including neatly aligned spermatid heads (Figure 5A-C). Sperm were also detected abundantly within the seminal vesicles (Figure 5D) and investment cones (marked by actin) were closely aligned with spermatid heads (Figure 5E). In testes from the *Hsc70Cb* RNAi 2 line (Figure 5G-I), spermatid bundles were not clearly visible and instead a build-up of entangled sperm was present at the distal testis (Figure 5G, H). In most cases no sperm were present in the seminal vesicles of the *Hsc70Cb* RNAi 2 line (Figure 5I). In rare instances, very few sperm were observed in the seminal vesicle (Figure 5J). Further, investment cones were disorganised and misaligned with sperm heads (Figure 5K). While even in the wild type setting, actin cones are not always associated with spermatid nuclei, this presentation was clearly abnormal.

**Figure 5.**
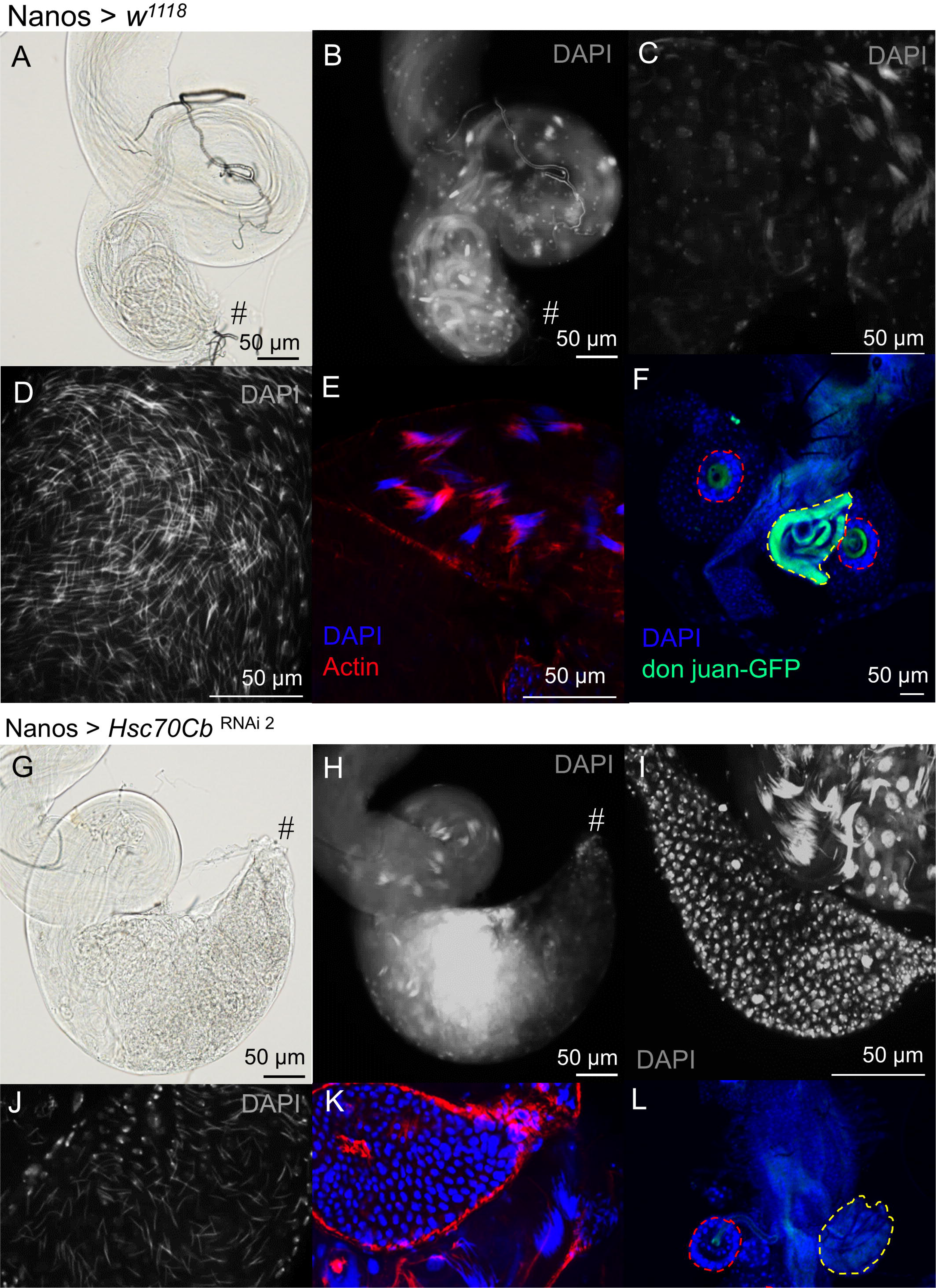
Sperm individualisation requires *Hsc70Cb* function. Testes and seminal vesicles were isolated from adult (A-E) control and (G-K) *Hsc70Cb* mutants (RNAi line 2) flies, stained with DAPI and probed for actin. (A, G) Brightfield and (B, H) DAPI stained images of the distal testis. The testis-seminal vesicle junction is denoted by #. (C, I) Spermatid individualisation within the distal testis and a connected seminal vesicle devoid of sperm in (I). (D, J) DAPI staining of seminal vesicles containing sperm, which is notably reduced in RNAi line 2. (E, K) Sperm individualisation complexes (actin, red). (F, L) Reproductive tracts of females mated with males from controls or RNAi line 2 mutants. Red circles trace the spermathecae and yellow circles trace the seminal receptacle, where green staining is indicative of the presence of sperm (tails).

We next mated wild type and mutant males containing don juan-GFP to *w^1118^* females, to test if sperm could be deposited in the female reproductive tract. When using control males, this approach revealed green fluorescence in both the spermathecae (red dashed lines) and notably within the seminal receptacle (yellow dashed line) (Figure 5F), indicating the presence of sperm in females. By contrast, no green fluorescence was seen in females mated by RNAi line 2 mutants, highlighting no sperm were present (Figure 5L). This analysis was not undertaken for RNAi line 1 in which no sperm were generated.

To determine the cellular content of tiny testes from mutant *Hsc70Cb* males resulting from RNAi line 1 knockdown in the germline, we next stained testis samples with markers of the hub (Fasciclin 3) and cyst stem and daughter cells (Broad) (Figure 6). In *w^1118^*control and RNAi line 2 males, Fasciclin 3 was characteristically localised to the ball of hub cells at the apical testis (Figure 6B, J). In RNAi line 1 males, however, Fasciclin 3 staining was observed throughout the testis and in an increased number of cells (Figure 6F). This is typical of testes that have lost germ cells, leading to an expansion of cells expressing hub markers (Gonczy and DiNardo, 1996). Similarly, Broad localisation was consistent in testes from both control and RNAi 2 line males: within cyst stem cells and early cyst cells (Figure 6D, L). By contrast, in testes from RNAi line 1 males, Broad positive cells, as a marker of cyst stem cells and early cyst cells, were localised further from the apical end of the testis, indicating that the somatic cell population had undergone aberrant proliferation (Gonczy and DiNardo, 1996). Collectively, these data suggest that the loss of *Hsc70Cb*, leads to a loss of the hub and somatic cell arrangement in the apical testis in RNA line 1 males. This phenotype commonly coincides with a loss of germline cells (Hetie et al., 2014).

**Figure 6.**
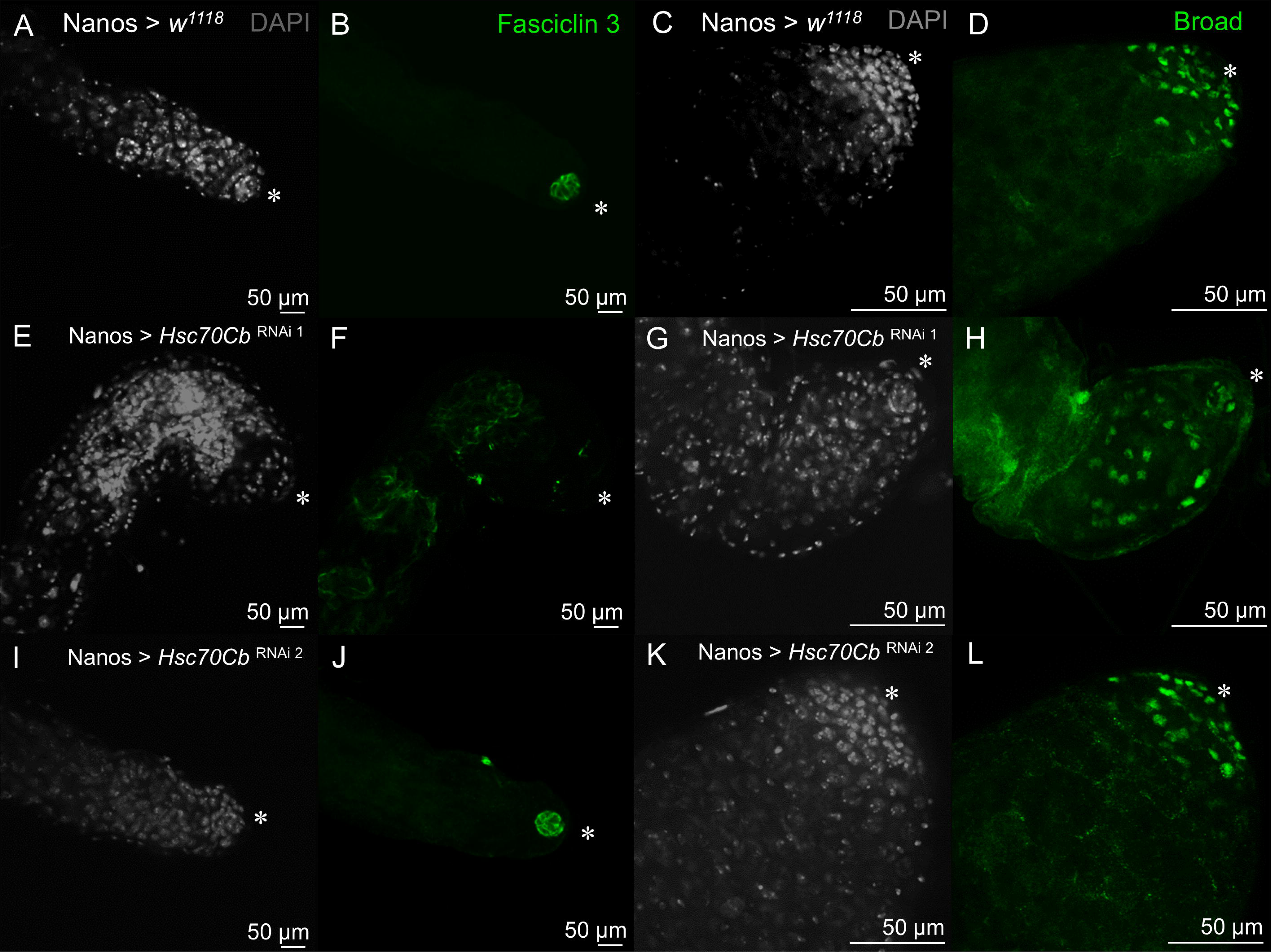
Organisation of the hub is altered in *Hsc70Cb* mutants. Adult testes were isolated from control and *Hsc70Cb* mutants and stained with Fasciclin *3* (hub marker) and Broad (cyst stem cells and progeny marker). (A-D) *w^1118^*, (E-H) *Hsc70Cb* RNAi line 1, (I-L) *Hsc70Cb* RNAi line 2. Scale bars = 50 µm, * = hub.

### *Drosophila Hsc70Cb* partly shares biological function(s) with human HSPA4 and HSPA4L cDNA

To test the degree of functional conservation between each of fly *Hsc70Cb* and human *HSPA4* and *HSPA4L*, we generated a compound stock for each RNAi line containing transgenic UAS-human cDNA for *HSPA4* or *HSPA4L*. Fertility assessment of male flies crossed to the Nanos-Gal4 driver revealed that the individual expression of human *HSPA4* or *HSPA4L* failed to rescue fertility in RNAi line 1 and 2 germline knockdowns as measured by mating (*p* < 0.0001; Figure 7A). Histological assessment of these males, however, revealed that expression of human *HSPA4* or *HSPA4L* could partially rescue the early germ cell loss infertility phenotype observed in RNAi line 1, i.e. a partial phenotypic rescue (Figure 7B-L). Specifically, when either HSPA4 or HSPA4L were expressed, spermatogonia and spermatocytes were present in RNAi line 1 mutant testes (Figure 4). Although spermatocytes were present, no spermatids were detected. The spermatocytes underwent cell death before spermiogenesis could occur and no round spermatids were present. *w^1118^* controls expressing *HSPA4* (Figure 7A, D) or *HSPA4L* (Figure 7G, J) had testes containing all germ cell types and sperm within the seminal vesicles. No noticeable rescue was observed when HSPA4 or HSPA4L was introduced into RNAi line 2. Accordingly, and as seen in RNAi line 2 alone (Figure 4), spermatids remained poorly individualised and failed to transit into the seminal vesicles. Due to lethality, it was not possible to combine all genetic elements to generate flies containing the RNAi construct and *Nanos* driver with both *HSPA4* and *HSPA4L*. As such, we could not test the capacity of a double *HSPA4* and *HSPA4L* to fully rescue male fertility in *Hsc70Cb* germline mutants.

**Figure 7.**
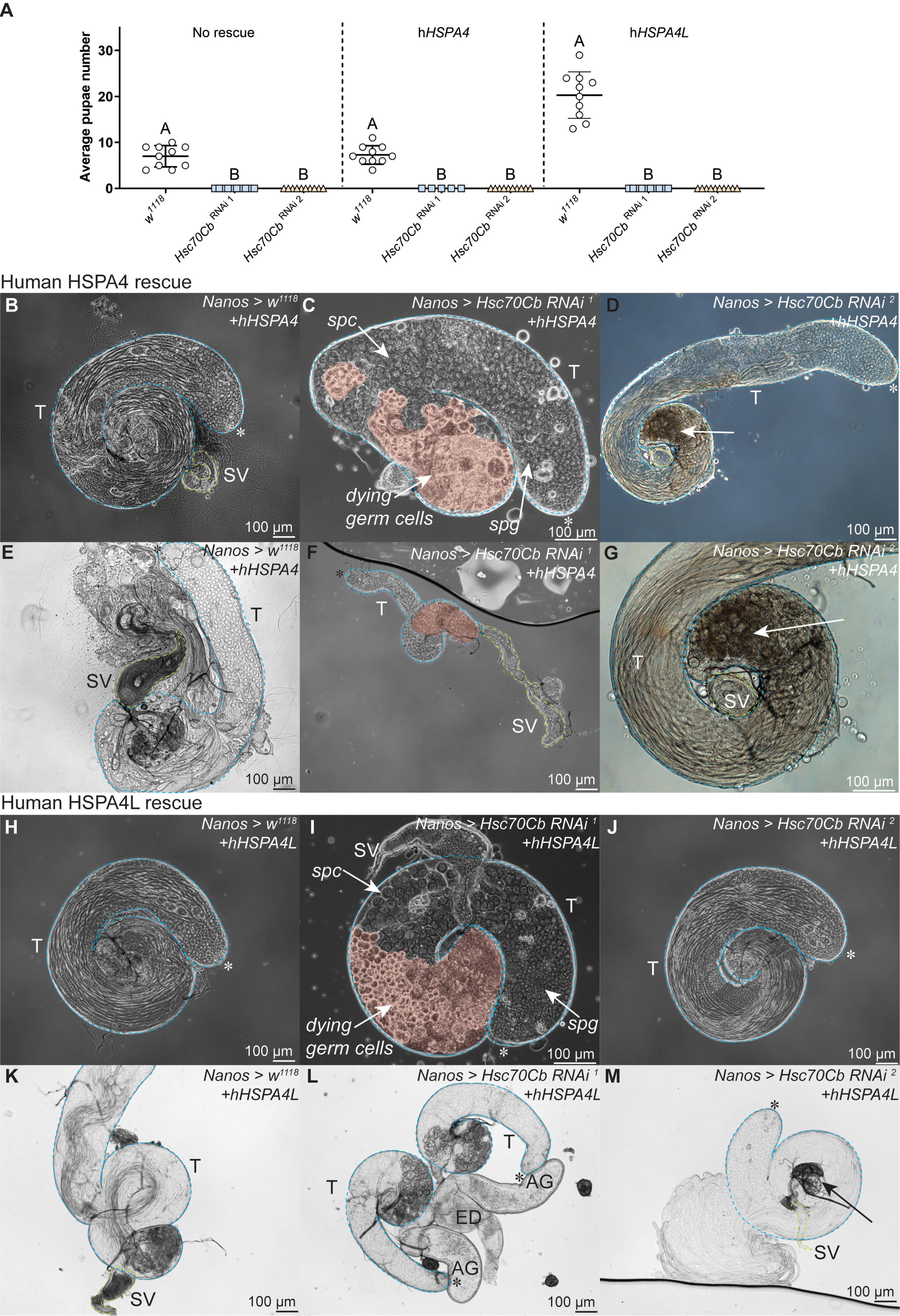
Human HSPA4 and HSPA4L partially rescue male germ cell loss in *Hsc70Cb* mutants. Fertility of *w^1118^* and *Hsc70Cb* RNAi 1 and 2 lines (driven by *Nanos*-Gal4), +/-co-expression of human (h) *HSPA4* or human *HSPA4L*, as measured by pupae counts. Adult testis histology from *w^1118^* and *Hsc70Cb* RNAi 1 and 2 lines containing hHSPA4 (B-G) or hHSPA4L (H-M). Arrows in D, G and M denote abnormally coiled sperm in the distal testis. Scale bars = 100 µm. T = testis, SV = seminal vesicle, AG = accessory gland, ED = ejaculatory duct, spg = spermatogonia, spc = spermatocyte, * = hub.

A similar partial rescue was observed for females in the *Hsc70Cb* RNAi line containing human HSPA4 and HSPAL (Supplementary Figure 1I-J, L-M), demonstrating the conserved role of *Hsc70Cb* in early germ cell development in both sexes.

In summary, these data highlight a shared gene function of the HSPA4 family in early germ cell development in both humans and flies. We define at least two roles of *Hsc70Cb* in male fertility, one in establishing germ cell development in spermatogonial stem cells that give rise to all germ cell types, and a second in regulating the process of spermatid individualisation.

## Discussion

Herein, we reveal the essential role of *Hsc70Cb* in multiple phases of *Drosophila* spermatogenesis. Aligning with single cell sequencing data, strong knockdown of *Hsc70Cb* in the male germline, as occurred in RNAi line 1, resulted in the absence of male germ cells and a tiny testis phenotype in both larvae and adults. A less robust knockdown of *Hsc70Cb,* with RNAi line 2, resulted in abnormal sperm individualisation, highlighting a requirement for *Hsc70Cb* in at least two phases of germ cell development to support male fertility – germline stem cell/spermatogonia survival and spermiogenesis. The absence of sperm in the *Hsc70Cb* RNAi line 1 mutants is consistent with the infertile man harbouring a homozygous *HSPA4L* genetic variant (Nagirnaja et al., 2022), supporting the hypothesis that *HSPA4L* is required for male fertility in humans and flies and highlighting the utility of the fly as a model of human spermatogenesis.

In flies a single orthologue, *Hsc70Cb*, exists, whereas in higher order species there has likely been gene duplication to produce two genes, *HSPA4* and *HSPA4L*. *Hsc70Cb* is also a predicted orthologue for a third heat shock protein in humans, *HSPH1*, highlighting potential neofunctionalisation of these heat shock proteins in higher order species. In this study, the loss of *Hsc70Cb* function in the fly germline resulted in male sterility, aligning with the human phenotype (Nagirnaja et al., 2022). By varying the degree of loss-of-function, we revealed two roles for Hsc70Cb: in spermatogonia function and sperm individualisation during spermiogenesis. Expression of either human *HSPA4* or *HSPA4L* in *Hsc70Cb* mutants partially rescued the germ cell loss phenotype, but infertility remains. Functionally, *Hsc70Cb* is predicted to act via its substrate binding-β subdomain to process unfolded polypeptides and prevent cytotoxic protein aggregation (Yakubu and Morano, 2021). To enact this role, Hsc70Cb interacts with a second heat shock protein, DNAJ-1 (*DNAJB1* in mammals) (Kuo et al., 2013). Additionally, *Hsc70Cb* may play a role in the prevention of amyloid formation via a C-terminal domain that is widely conserved across metazoans (Kuo et al., 2013, Yakubu and Morano, 2021). Our results suggest a crucial role for *Hsc70Cb* in the prevention of protein aggregation in male germ cells. This may suggest that additional heat shock protein function, potentially as a chaperone, or the combined effects of HSPA4 and HSPA4L are required to restore fertility in *Hsc70Cb* germline mutants.

In mice, the deletion of either *Hspa4* or *Hspa4l* caused infertility in a subset of males (Held et al., 2011, Held et al., 2006). For those *Hspa4*^-/-^ male males that were fertile (42%), they produced fewer sperm than wild type males. Similarly, while a portion of *Hspa4l*^-/-^ male mice could sire offspring (39%), litters were smaller than wild type. The deletion of both *Hspa4* and *Hspa4l* in mice resulted in neonatal lethality due to compromised lung function arising from impaired chaperone activity and elevated protein ubiquitination (Mohamed et al., 2014). When comparing *Hspa4l*^-/-^ mice to humans, the phenotype is mostly concordant, with reduced sperm count and subfertility (Held et al., 2006). It is possible that the presence of environmental (e.g. heat stress) and/or lifestyle effects is required in mice to exacerbate the infertility presentation to that similar of the azoospermia man carrying a *HSPA4L* variant. Indeed, heat shock proteins were discovered in *Drosophila* as a response to cellular heat stress (Tissieres et al., 1974).

Collectively, these data highlight the crucial role of the *HSPA4* gene family in male fertility across animals broadly. The localisation of HSPA4L to the midpiece in sperm and the association of HSPA4L content with sperm motility in humans, highlight a conserved role of HSPA4L in sperm formation and function across mammals. Several other heat shock proteins are also expressed throughout germ cell development and are required for sperm function (Nixon et al., 2017, Ji et al., 2012), noting that spermatogenesis is exquisitely sensitive to heat stress (Houston et al., 2018). For example, *hsp60B* is similarly essential for sperm individualisation in *Drosophila* (Timakov and Zhang, 2001) and *Hsp90*β*1* is required for sperm head morphogenesis in mice (Audouard and Christians, 2011). Specifically, *hsp60B* loss resulted in poor function of the individualisation complex, similar to the individualisation defects in the *Hsc70Cb* mutant (RNA line 2) in this study. Outer dense fibre protein 1, which is also classified a small heat shock protein (HSPB10), plays essential roles in sperm tail development and head-tail coupling apparatus function (Yang et al., 2012). While other heat shock proteins are not essential for fertility, they are involved in germ cell homeostasis through the prevention of oxidative stress and the regulation of apoptosis in germ cells, optimising sperm quality (Purandhar et al., 2014).

In summary, in this study we reveal the conserved role of the HSPA4 gene family in male fertility across mammals and provide evidence that variants in the *HSPA4L* gene cause infertility in flies and men.

## Author Roles

BJH, JN, RB and ANA undertook experiments

BJH, RB, GH and MKOB wrote the paper

BJH, RB, GH and MKOB provided the funding and administration

## Figure Legends

**Supplementary Figure 1. *Hsc70Cb* is essential for oocyte development.** (A) Fertility of female control and *Hsc70Cb* mutants. (B-G) Adult ovaries were isolated from control and *Hsc70Cb* mutants and stained with DAPI. (B, E) Brightfield images of control and *Hsc70Cb* RNAi line mutant ovaries. DAPI stained ovaries from control (C-D) and *Hsc70Cb* RNAi line 1 mutants (F-G), with a representative ovariole highlighted in white dashed lines. Phase contrast imaging was used to visualise histology of ovaries from control, mutant and HSPA4L rescue flies (H-M). (H) Control, (I-J) *Hsc70Cb* RNAi line 1, (K) Control with HSPA4L, (L-M) *Hsc70Cb* RNAi line 1 with HSPA4L. * = mature eggs, arrows = partially rescued egg development.

**Supplementary Figure 2. Mutant testis histology.** Testes were isolated from adult control and mutant *Hsc70Cb* flies and stained with DAPI. (A) *w^1118^*control testis morphology. (B) Tiny testes and seminal vesicles in *Hsc70Cb* germ cell knockdown RNAi line 1. (C) Spermatid aggregation (arrow) in *Hsc70Cb* germ cell knockdown RNAi line 2 (arrow). Scale bars = 50 µm. T = testis, SV = seminal vesicle, ED = ejaculatory duct, * = hub.

