## Supplementary figures and images for "Mammalian heat shock protein A4 family ortholog *Hsc70Cb* is required for two phases of spermatogenesis in *D. melanogaster*"

### Supplementary Figure 1

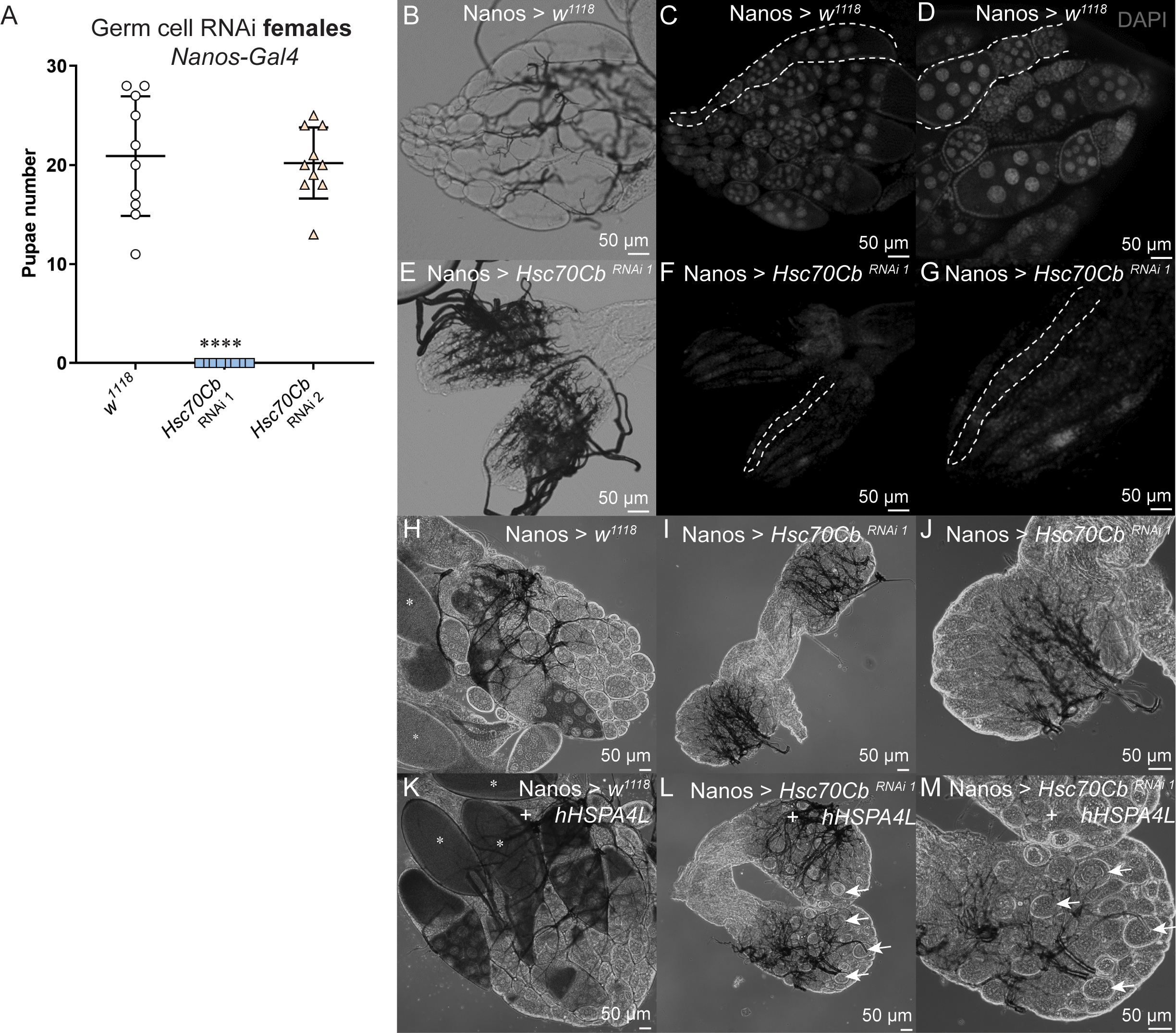

### Supplementary Figure 2

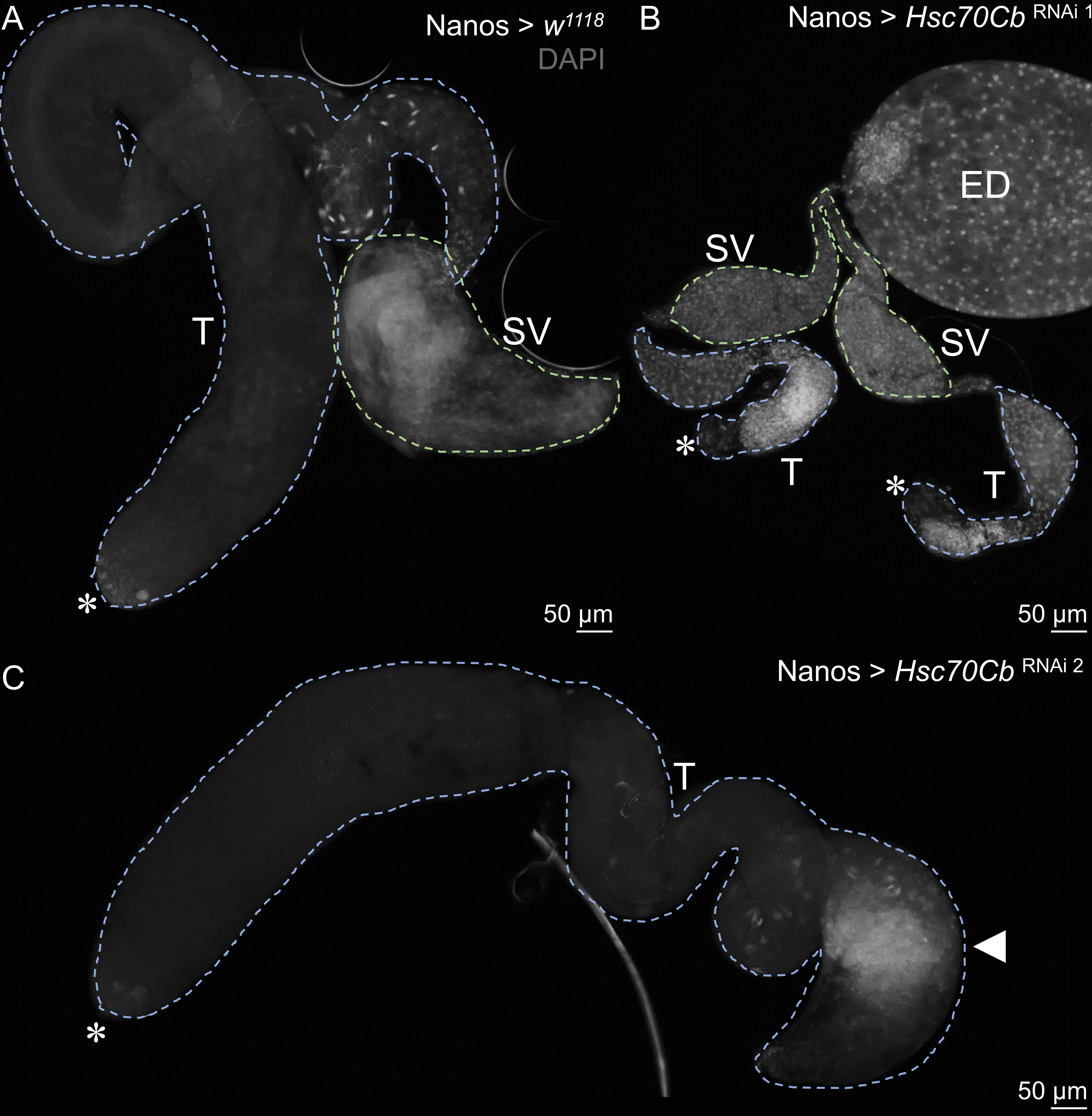
